# Mechanistic modeling of cell viability assays with *in silico* lineage tracing

**DOI:** 10.1101/2024.08.23.609433

**Authors:** Arnab Mutsuddy, Jonah R. Huggins, Aurore K. Amrit, Atalanta Manuela Harley-Gasaway, Cemal Erdem, Jon C. Calhoun, Marc R. Birtwistle

## Abstract

Data from cell viability assays, which measure cumulative division and death events in a population and reflect substantial cellular heterogeneity, are widely available. However, interpreting such data with mechanistic computational models is hindered because direct model/data comparison is often muddled. We developed an algorithm that tracks simulated division and death events in mechanistically-detailed single-cell lineages to enable such a model/data comparison and suggest causes of cell-cell drug response variability. Using our previously developed model of mammalian single-cell proliferation and death signaling, we simulated drug dose response experiments for four targeted anti-cancer drugs (alpelisib, neratinib, trametinib and palbociclib) and compared them to experimental data. Simulations are consistent with data for strong growth inhibition by trametinib (MEK inhibitor) and overall lack of efficacy for alpelisib (PI-3K inhibitor), but are inconsistent with data for palbociclib (CDK4/6 inhibitor) and neratinib (EGFR inhibitor). Model/data inconsistencies suggest that (i) the importance of CDK4/6 for driving the cell cycle may be overestimated, and (ii) the cellular balance between basal (tonic) and ligand-induced signaling is a critical determinant of receptor inhibitor response. Simulations show subpopulations of rapidly and slowly dividing cells in both control and drug-treated conditions. Variations in mother cells prior to drug treatment impinging on ERK pathway activity are associated with the rapidly dividing phenotype and trametinib resistance. This work lays a foundation for the application of mechanistic modeling to large-scale cell viability datasets and better understanding determinants of cellular heterogeneity in drug response.

**Author Summary:** There is a large amount of data in the public domain for how a variety of cancer cell types respond to a multitude of anti-cancer drugs. However, interpreting such data mechanistically is hindered because of a lack of precise computational tools for model/data comparison. We developed an algorithm that tracks simulated division and death events in mechanistically detailed single-cell lineages to enable such a model/data comparison and suggest causes of drug response variability. We applied this tool to understand four targeted anti-cancer drugs better. Simulations are consistent with data for strong growth inhibition by trametinib and overall lack of efficacy for alpelisib, but are inconsistent with data for palbociclib and neratinib. Model/data inconsistencies suggest where biological understanding may be incomplete. Simulations show subpopulations of rapidly and slowly dividing cells in both control and drug-treated conditions, and analysis thereof suggest mechanisms potentially involved with acute drug resistance. This work lays a foundation for using large-scale datasets to better understand drug response.

## Introduction

One of the grand challenges of systems biology is to build a comprehensive and quantitative understanding of the structure and functionality of living cells. Whole cell models that describe the function of every gene and its products is an attractive manifestation of such a goal. Whole-cell models for some microorganisms exist^1–4^, but have not been reported for human cells. Nevertheless, a wide range of individual pathway^5–20^ and integrative multiple pathway^21–28^ human cell models have been published, which are a stepping stone^29,30^. These models can contribute to better understanding multiscale phenotypes^31–35^, diagnosis of disease states and their progression^36,37^, and development of efficacious therapeutic procedures^38–40^.

Data availability is a primary challenge for human whole cell modeling^29,41,42^. Large-scale databases^43–45^ containing cell viability assay data exploring sensitivity of multiple cell lines to multiple drugs are an attractive resource in this regard. Assessing how well computational models, based on current knowledge, explain this data can reveal existing knowledge gaps and inform next stages of research.

An obvious pre-requisite to leveraging such data sets for modeling purposes is the existence of robust methods for comparing simulations to experimental readouts appropriately. Most viability assays measure cell population size or a proxy. Typical mechanistic models do not explicitly describe cell division and death events, whose balance dictates cell population size, but usually prescribe empirical relations^22,46^. Course-grained agent based modeling^47–49^ can account for division and death events and were described in, for example, process control and optimization of production of therapeutic proteins using mammalian cell culture^50^, analyzing the impact of cell population heterogeneity in colony and tissue context^51^, and elucidating the role of heterogeneity in IFNβ signaling^52^. However, there remains a need for algorithms that can simultaneously track large-scale mechanistic detail of drug response with cell division/death events. Such algorithms would help interpret the large body of cell viability response data for building better human whole-cell models.

In this work, we present an algorithm that combines detailed mechanistic descriptions of drug action with single cell lineage-resolved division and death events to construct simulation outputs that are directly comparable to cell viability assay data. As a test case, we use a previously-developed, large-scale model of single mammalian breast epithelial cell (MCF10A) proliferation and death, SPARCED^53^, and add to it mechanistic pharmacodynamic models based on known binding interactions between drugs and modeled targets. First, we describe the algorithm and the types of novel analytics that can be derived, such as cell population size dynamics, cross-generational biomarker tracking for cell lineages, and cell population dendrograms. Then, we simulate dose responses to multiple drugs for which experimental data exist, namely, alpelisib (PI-3K inhibitor), trametinib (MEK inhibitor), palbociclib (CDK4/6 inhibitor) and neratinib (EGFR inhibitor). Simulations agree with experiments for strong growth inhibition by trametinib and overall lack of efficacy of alpelisib; however, there is substantial model-experiment discrepancy for palbociclib and neratinib. Analysis of model discrepancies suggests that (i) the importance of CDK4/6 for driving the cell cycle is likely overestimated, and (ii) the cellular balance between basal (tonic) and ligand-induced ERK signaling is a critical determinant of EGFR inhibitor response. We also applied the model to better understand an interesting phenomena that simulations showed subpopulations of rapidly and slowly dividing cells in both control and drug-treated conditions. We find that variations in mother cells prior to drug treatment impinging on ERK pathway activity are associated with the rapidly dividing phenotype and trametinib responsiveness. This work establishes a foundational framework for applying mechanistic modeling to large-scale cell viability datasets, which are essential for developing comprehensive human cell models. Additionally, it offers a unique analytical approach to generate hypotheses regarding the molecular drivers of cellular variability across generations.

## Results

### Lineage-resolved single-cell simulation framework

We first set out to construct a simulation algorithm that mirrors drug dose response viability assays (**Fig. 1**). These assays typically start with a population of asynchronously cycling cells that are treated with drug for ∼3 days and then assayed for final cell number (or a metric proportional to it). The final cell number is related to the number of division and death events each initial single cell ultimately experienced. Thus, the simulation algorithm starts with a population of asynchronously cycling cells and counts the individual division and death events from each initial cell. Throughout this manuscript, we use our previously published mechanistic model (SPARCED) of single cell proliferation and death signaling^53^ representing MCF10A breast epithelial cells. In principle, however, any model with single-cell resolution and division/death readouts is potentially compatible with the approach. We provide one such simple example^54^ in the Methods and GitHub repository.

**Figure 1.**
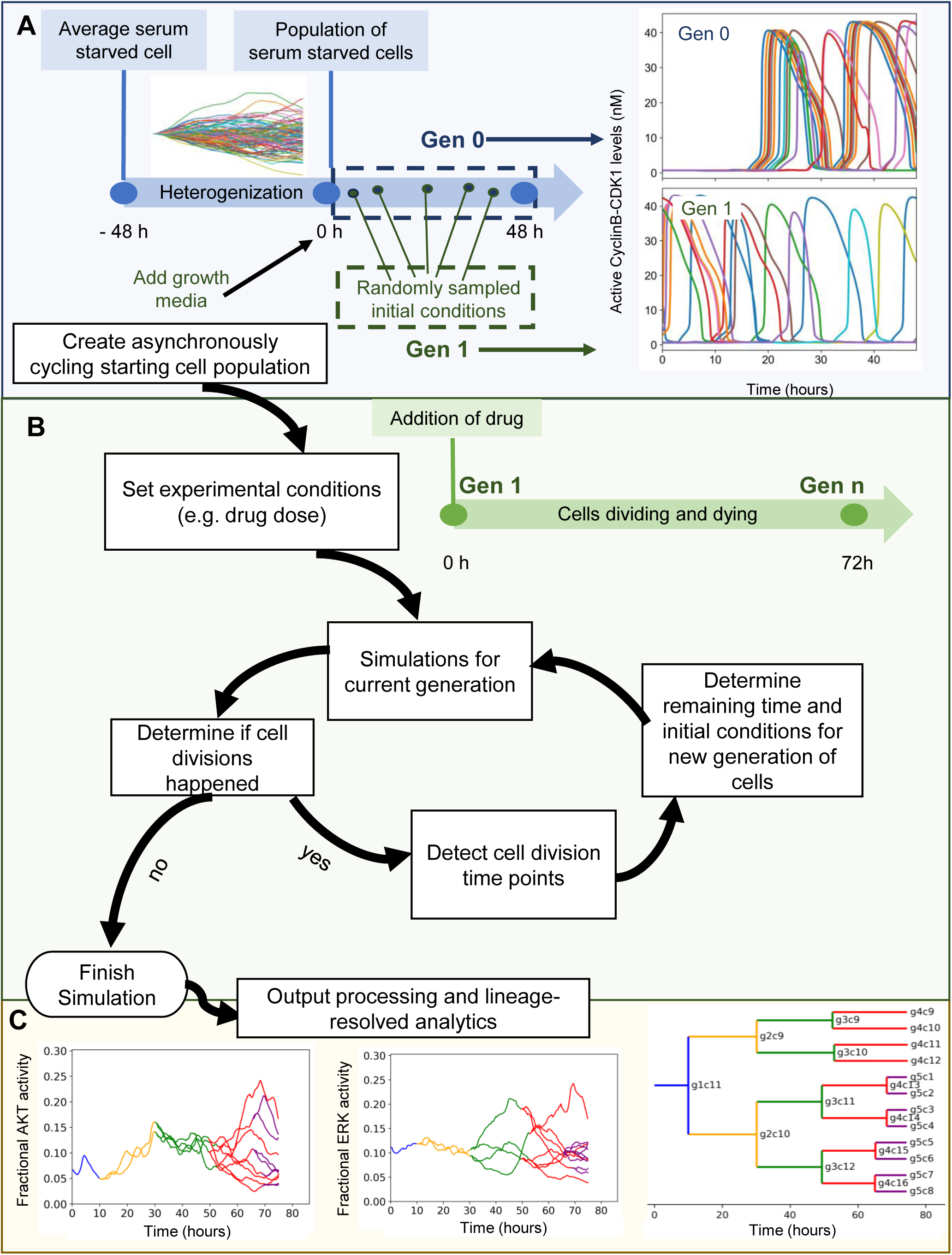
Workflow of the developed simulation algorithm. **A.** An asynchronously cycling cell population (Gen 1) is initiated by sampling the initial conditions at random time points from a pool of single cell simulations run with growth factor stimulation (Gen 0). **B.** Upon the execution of each generation, detection of new cell division events (or lack thereof) within simulation time determines the creation or not of a next generation. **C.** (left and center) Cross-generational trajectory of observed ERK and AKT activity from a randomly-chosen single-cell lineage. Varying colors represent subsequent generations, starting from Gen 1. (right) *In silico* lineag1e tracing capability is demonstrated with a lineage dendrogram. Lines representing individual cells are labeled with generation and index.

The algorithm begins with the creation of a simulated population of asynchronously-cycling, single cells (**Fig. 1A**). The initial model state is an average, serum-starved cell (non-cycling). The first step is to generate a population of cells with heterogeneous gene expression profiles, enabled by descriptions of intrinsic noise in gene expression as previously described^21^. We refer to this process as “heterogenization”. After 48 simulated hours, when the distribution of most protein levels across the cell population stabilizes, the addition of growth media is simulated (in the case of MCF10A cells and this model—EGF and insulin). Subsequently, synchronized cell cycle progression is observed in simulations for an additional 48 hours, creating so-called “Generation 0” (Gen 0). To convert these Gen 0 synchronized cells into Gen 1 asynchronously cycling cells, we sample random times from the 48 hour growth media treatment window for each single cell. These selections become initial conditions for Gen 1, which is then subjected to simulated drug treatment for 72 hours (**Fig. 1B**).

Once these simulations are completed, the outputs are analyzed to determine cell division events (based on Cyclin B-CDK1 peaks for this model) and their time points (**Fig. 1B**). Based on the cell division time points, the remaining simulation time (difference between division time and 72 hours), and initial conditions for each daughter cell are determined for the next generation, and lineage information is recorded. Importantly, in SPARCED, daughter cells immediately begin experiencing drift from one another due to stochastic gene expression, which is constantly occurring in every simulated cell differently. Subsequently, simulations for the next generation are run and this cycle continues until no division events occur in a given generation. Detected cell death events (based on cleaved PARP dynamics for this model), halt a lineage.

These simulations not only mirror typical drug dose response experiments but also enable lineage-resolved analyses (**Fig. 1C**). Since individual division and death events from a parental cell are tracked, it also allows dynamic tracking of observables (such as ERK or Akt activity) across multiple generations of any single cell lineage. Lineage dendrograms can be also constructed, as is typical in such analyses. Such capability may generate hypotheses linking drug sensitivity or resistance with cell fates and lineage, or variations in biochemistry that predispose cells to response or resistance.

### Comparing Simulated Drug Dose Responses to Experimental Measurements

With the above algorithm, we could now compare model predictions to experimental data for cell viability drug responses. Specifically, we focused on four previously-studied, targeted anti-cancer drugs for which our model includes primary targets and significant off-targets: trametinib (MEK inhibitor), alpelisib (PI-3Kα inhibitor), neratinib (EGFR/ErbB2/ErbB4 inhibitor), and palbociclib (CDK4/6 inhibitor)^55^. We extended SPARCED by including known drug interactions with target species^56–59^ and leveraging previously described capabilities to robustly and easily increase the model scope^53^ (see Methods).

We then performed lineage-resolved simulations for various doses of the modeled drugs for which experimental data are available^55^, with no adjustment to the SPARCED model (besides addition of the drug pharmacodynamic modules—see Methods), using a starting population of 100 cells. This framework allows direct simulation of the dynamic cell population size in response to drug doses (**Fig. 2A**). The effect of drug action can be visualized by cell lineage dendrograms, showing in this example the clear effect of moderate trametinib doses to reduce the number of cell division events (**Fig. 2B-C**). The simulation outputs were used to calculate growth-rate inhibition^60^ metrics which is the same method applied to the experimental dataset, allowing direct comparison of experimental and simulation results (**Fig. 2D-G**).

**Figure 2.**
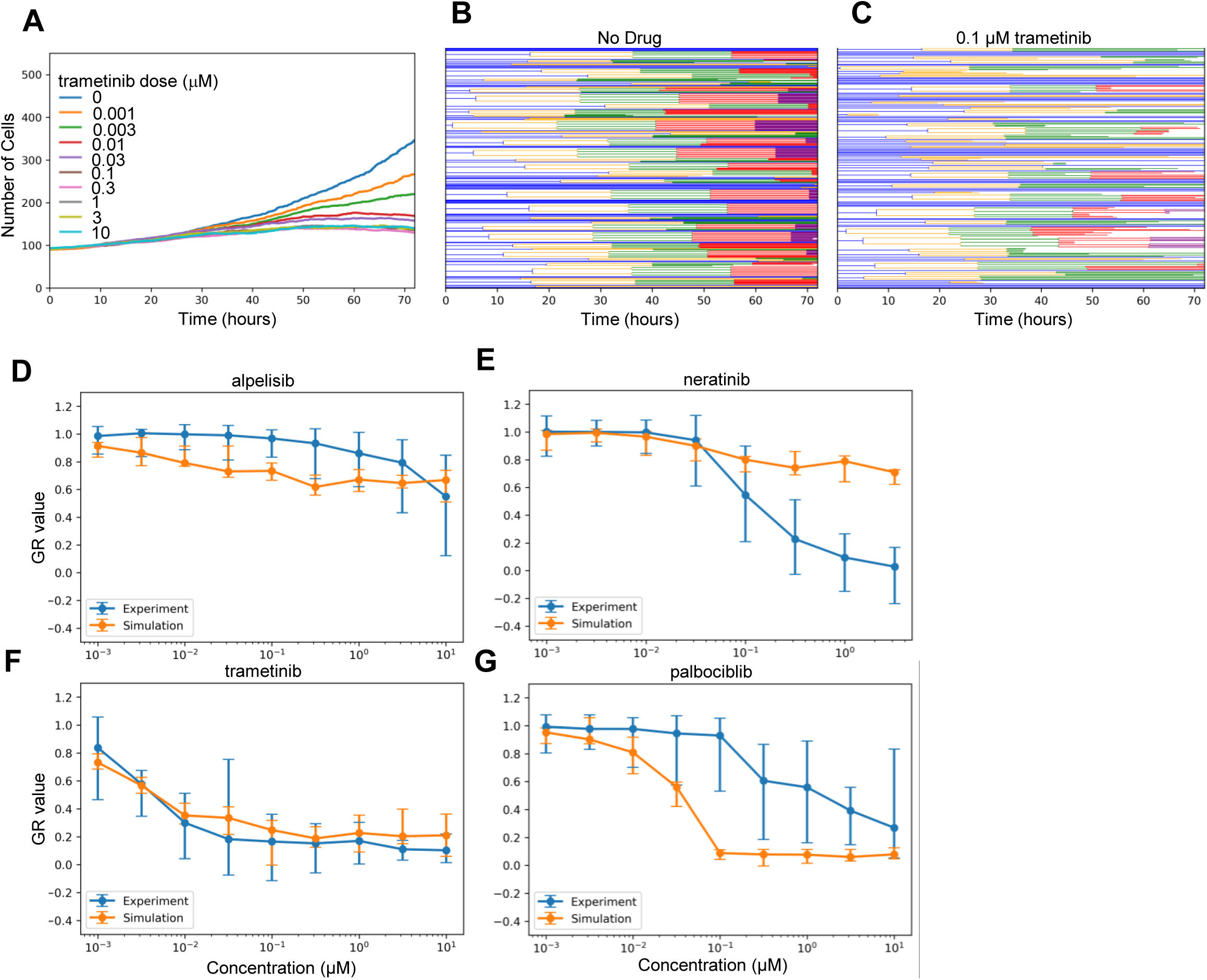
Lineage Resolved Simulations for Comparing Simulated and Experimental Cell Viability Assays. **A-C.** Dose response simulations for an example drug (trametinib). Median (across simulation replicates) cell population dynamics for several doses (A) and cell population lineage dendrograms for specific doses: 0 µM (B) and 0.1 µM (C) are shown. **D-G.** Simulated dose responses measured in GR-value for four drugs compared experimental data. Error bars are standard error taken from original experimental data, or as calculated across simulation replicates (n=10).

The simulation results for trametinib (**Fig. 2F**) demonstrate surprising agreement with experimental data considered no parameter fitting was done, and also expectedly indicate substantial cell-to-cell variability at low doses. The simulations also captured the overall lack of efficacy for alpelisib, albeit with some slight differences in dose response slope (**Fig. 2D**). Simplified representations of PI-3K biology in the underlying model, which does not account for isoform-specific effects, could explain part of the difference. Alpelisib is a PI-3Kα isoform-specific inhibitor, but is modeled necessarily as a pan-PI-3K inhibitor. Comparison to data for other inhibitors such as pictilisib, a pan-PI-3K inhibitor, and taselisib, a beta-sparing inhibitor, could help constrain such efforts, although the model would need to be expanded to account for the increased and unmodeled off-targets of these drugs relative to alpelisib^61^.

On the other hand, predicted palbociclib (**Fig. 2G**) and neratinib (**Fig. 2E**) responses were substantially different from experiments. Subsequent analyses explore the nature of these differences, as well as potential reasons for the cell-to-cell variability in the trametinib response. We also note that independent cell death data were available^61,62^, which largely showed these drugs do not substantially kill cells, roughly consistent with GR values greater than 0, and with simulations (**Fig. S1**). A partial exception is neratinib, which shows some cell death at high doses, which may be due to general toxicity because it is an irreversible inhibitor.

### Palbociclib Dose Response Discrepancies Suggests CDK4/6 is Partially Redundant for Cell Cycle Progression

What could explain the experiment/simulation discrepancy for palbociclib, a potent inhibitor of CDK4/6, canonically understood to be a central mediator of cell cycle progression from G_0_ and G_1_ to S-phase^63–65^ (**Fig. 2G**)? Simulated palbociclib dose response starts to deviate from the experimental results at doses as low as 0.01 μM. Above 0.1 μM, the simulated dose response shows complete cytostasis. On the other hand, experimental results show minimal growth inhibition at 0.1 μM, and even high doses indicate only partial growth inhibition.

One consideration was doubling time. We reasoned that if the experimental doubling time was slower (greater) than the simulated doubling time, then in simulations many more cell divisions would be inhibited by the drug. That may explain why the simulated effects of palbociclib were much greater than that observed in experiments. The used GR metric in principle should help to account for such doubling time-related phenomena^60^, but it also relies on assumptions such as exponential growth and constant drug effect on growth, which may not be satisfied. The model predicted a slower doubling time (∼48 hours) than was reported in experiments for these MCF10A cells (∼18-25 hours) (**Fig. 3A** and ^55^), although a wide range is reported for this cell line (∼48^66^ or even ∼96 hrs^67^). This is opposite of the difference we expected may explain the dose response curve discrepancy. Therefore, we conclude doubling time differences are unlikely to explain the observed differences.

**Figure 3.**
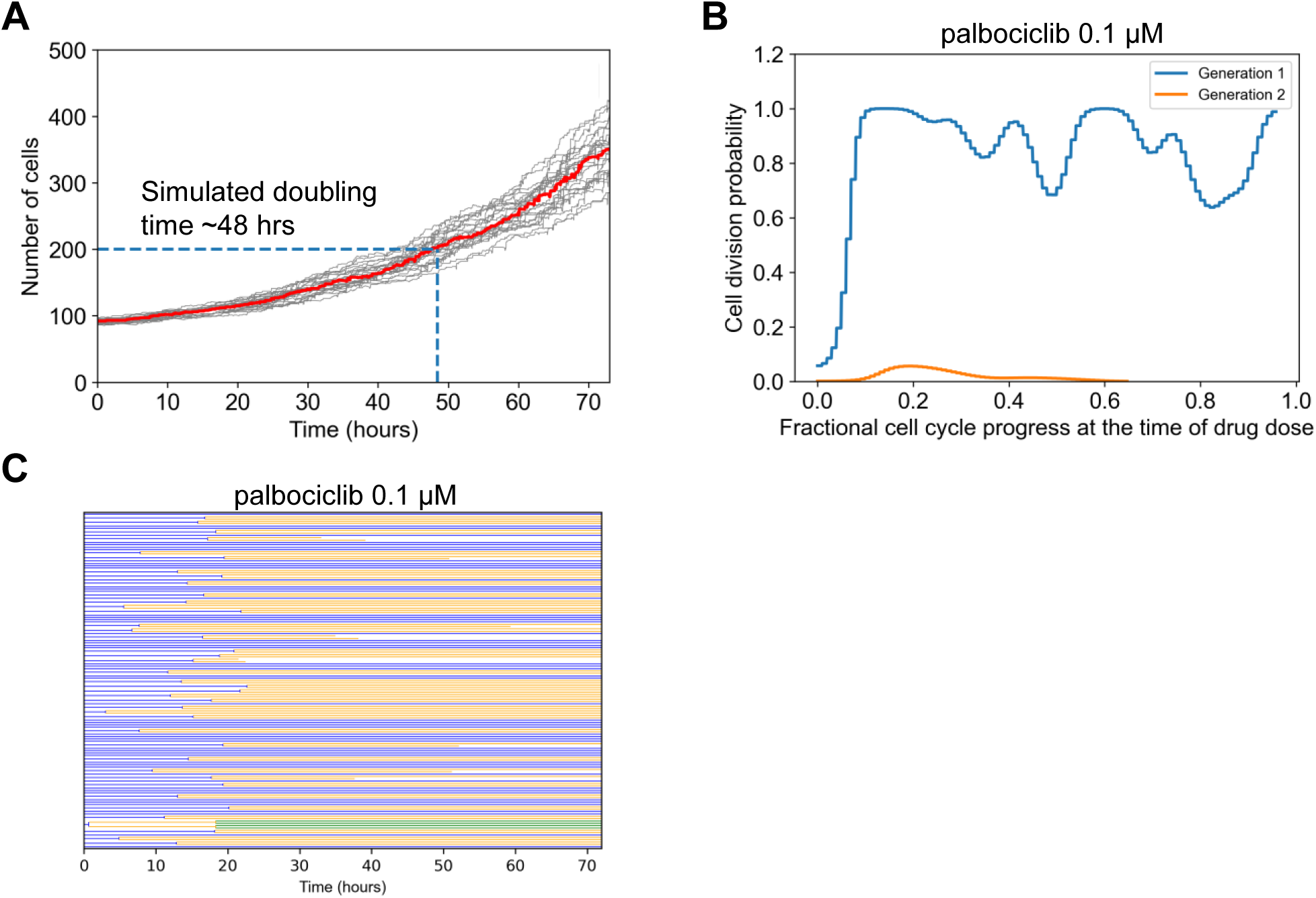
Simulation Analysis to Investigate Palbociclib Dose Response Discrepancy. **A.** Simulated cell growth curves under control conditions for multiple replicates. The dark black line is the median which was used to estimate doubling time, when the initial cell number (100) doubled (200). **B.** The fractional progression through the cell cycle (see Methods) was estimated for the beginning 100 generation 1 cells. This was the point at which 0.1 μM palbociclib was administered. The division outcome for each cell was then determined for this current generation, and if it exists for the next generation. The probability of division occurring was empirically estimated from this collection of binary outcomes and then plotted. **C.** Cell lineage dendrogram for response to 0.1 μM palbociclib. Most cells divide once early, and then the response is cytostatic.

Another consideration was restriction point behavior with respect to CDK4/6 activity, where a cell continues to divide even after complete inhibition of CDK4/6 at some point in the cell cycle^68–70^. We reasoned that if the model does not capture such behavior, simulated palbociclib treatment could immediately stop cell divisions instead of letting already committed cells continue to divide once, leading to more predicted potency than observed. To explore this in simulations, we analyzed the probability of cell division in Gen 1 versus Gen 2 cells, as a function of dynamic progress in the cell cycle at the time of simulated drug treatment (**Fig. 3B**). Prior studies place the restriction point early in the cell cycle^68^. Simulations with saturating palbociclib dose (0.1 μM) reflect such behavior, where most cells divide once if the cell cycle is at least ∼10% completed, but subsequent cell divisions are nearly non-existent. This effect is also clear from the simulated lineage dendrogram which shows most cells divide once but not subsequently with this dose of palbociclib (**Fig. 3C**). Thus, we conclude that modeled restriction point behavior is also unlikely to explain discrepancies.

Finally, we considered that the canonically understood role of CDK4/6 as modeled in SPARCED is simply inadequate. That is, the assertion that CDK4/6 activity is a necessary and sufficient step to drive the early cell cycle may be inaccurate^68,71,72^. A clinical line of evidence is the fact that CDK4/6 inhibitors have limited efficacy outside of hormone-positive breast cancers^63^. It has also been reported that proliferation can occur in CDK4/6 knockout cells^73^. More recent data have suggested that CDK4/6 activity has more of a probabilistic effect on cell cycle progression^74^, and the restriction point may be more reversible than previously thought in response to CDK4/6 inhibition^75^. CDK activities may also be overlapping; for example CDK2 and CDK4/6 may be compensatory^76^, and a sensor integrating multiple CDK activities^77^ was shown to be highly predictive of restriction point behavior^78^. Therefore, we conclude that most likely, fundamental model reformulation is needed to capture the effects of palbociclib, and that the canonical view of CDK4/6 as necessary and sufficient for cell cycle progression may be inadequate.

### The Balance of Tonic Versus Ligand-Induced Growth Factor Signaling is Critical for Capturing Drug Effects

Neratinib is an irreversible inhibitor of the EGFR (with some off-target activity for the closely related ErbB2/HER2 and ErbB4/HER4), a receptor tyrosine kinase that, upon ligand binding, activates the pro-proliferative and -survival ERK and AKT pathways^79–81^. Hence, drug action is expected to block ERK and AKT signaling when a ligand, such as EGF, binds to EGFR. The experimental dose response (**Fig. 2E**) shows strong growth inhibition at doses above 0.1 μM and complete cytostasis at ∼3 μM. However, simulation-predicted growth inhibition within this range is significantly weaker.

To explain this discrepancy, we considered that the current modeled balance of ligand-induced versus basal (also called tonic) ERK signaling could be incorrect. Specifically, that basal ERK signaling was too strong and causes non-negligible proliferation in the absence of EGF. If cell cycling is initiated by basal signaling too strongly, coupled with the fact that neratinib cannot inhibit basal signaling, this could explain some of the model-experiment discrepancy.

MCF10A cells are dependent upon EGF for cell cycle progression^82,83^. Thus, in simulations, cells dividing without EGF would support the above explanation. In simulations where the growth media contained only insulin, some cell division events were observed (**Fig. 4A**). Since the proliferative signaling activity that caused these divisions did not originate as a result of simulated EGF-EGFR activity, simulated neratinib treatment cannot inhibit these. This is inconsistent with the experimentally observed cell behavior and hence may be a major cause of mismatch between simulation and experiment.

**Figure 4.**
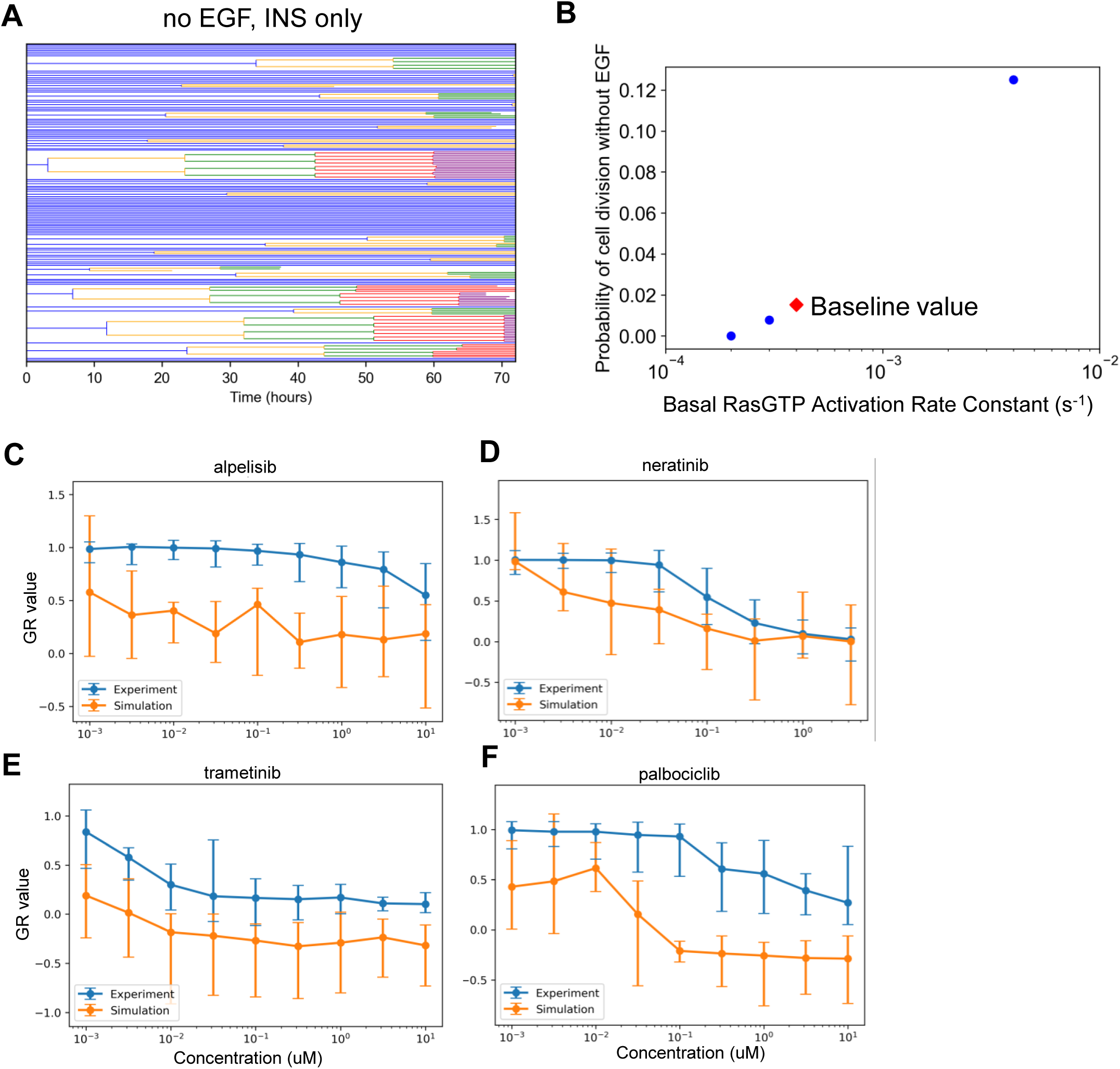
Simulation Analysis to Investigate Neratinib Dose Response Discrepancy. **A.** Lineage dendrograms under control (no drug) conditions without EGF but with insulin. There are multiple rapidly dividing cells. **B.** Dependence of the probability of cell division as a function of the rate constant controlling basal Ras activation. Simulations were done as in A, without EGF and with insulin. The baseline value for the rate constant in the current published version of the model is designated by the red diamond. **C-F.** Dose response curves as in Figure 2, except with the altered basal RasGTP activation rate constant from panel B (2x10^-4^ s^-1^).

How could the model be changed to account for these mismatches? First, we ensured that basal ERK signaling in the presence of insulin minimally induces cell cycle progression. Basal Ras-GDP to Ras-GTP exchange is the main reaction controlling basal ERK activity in the model. We reduced the value of the associated rate constant until the probability of cell division in the absence of EGF and presence of insulin was near zero (**Fig. 4B**—last point on left, 2x10^-4^ s^-1^), and then simulated the dose responses again (**Fig. 4C-F**). The new simulated neratinib dose responses show closer alignment with experiments. However, for all other drugs, experiment-model agreement became significantly worse, most likely now because the absolute levels of EGF-induced ERK signaling are altered. This result reinforces the close interacting nature of signaling mechanisms in the model for influencing broad features of drug response, and cautions against developing models without considering comparison to a compendium of data. Further model refinement in this regard, therefore, will be the scope of future work.

### Explaining Single-Cell Heterogeneity in Division Rate and Trametinib Response

A commonly observed phenotypic variation among cells within a population is the division rate^84,85^. Under both control (no drug) and trametinib (∼half-maximal response dose = 0.03 nM) treatment conditions, simulations show large variability in the number of divisions arising from a particular Gen 1 mother cell (**Figs. 5A-B**). Since rapidly dividing simulated cells (indicated by red) are present prior to drug treatment and persist after drug treatment, in simulations they are largely responsible for the partial response to trametinib. Could properties in the initial state of Gen 1 mother cells at the time of trametinib treatment be a predictor of resistance, in this case marked by persistent rapid division in the presence of drug? Do the same mechanisms that drive rapid division in control conditions apply to this drug resistance?

**Figure 5.**
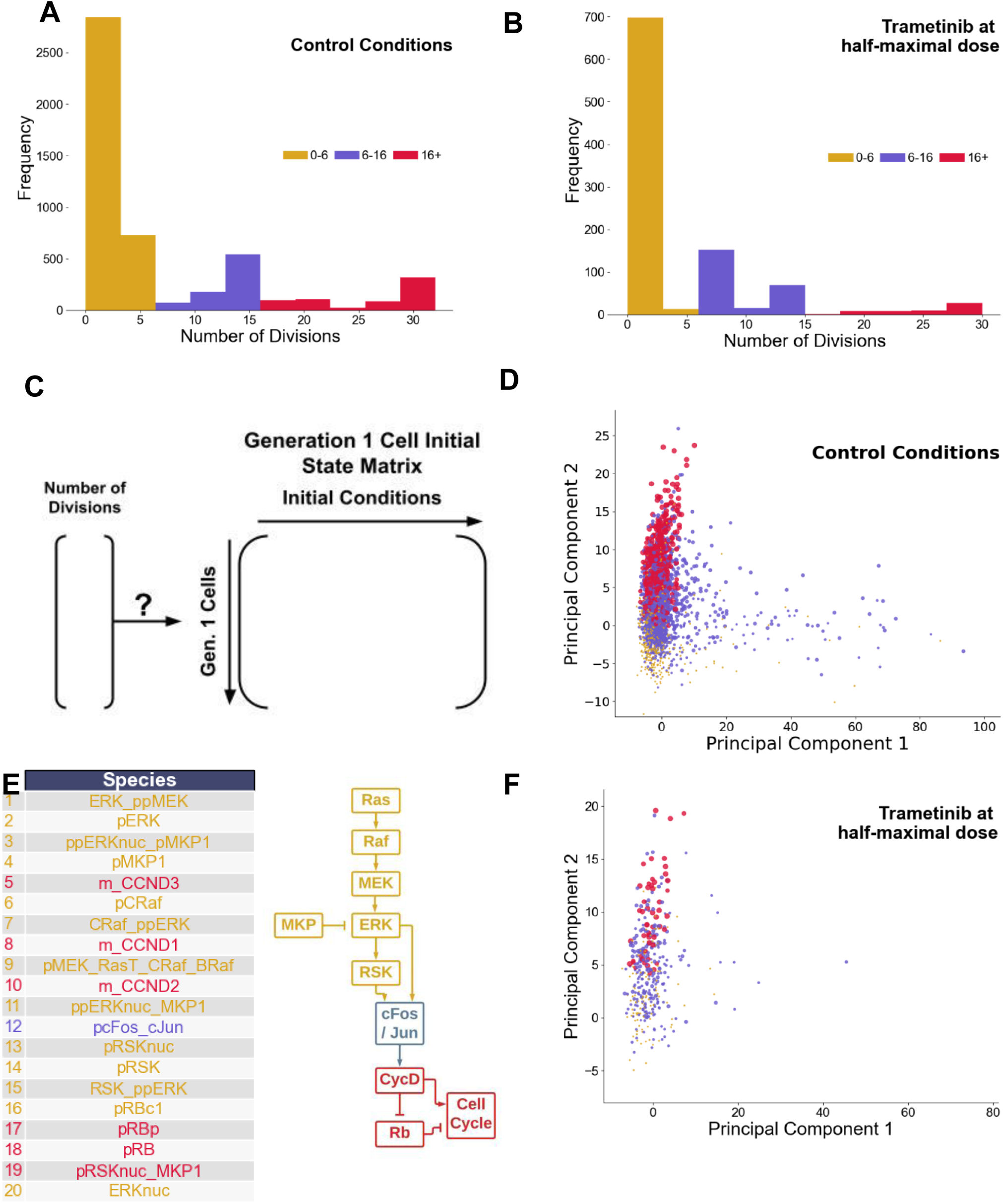
Rapidly Dividing Phenotype in Control and Trametinib-Treated Conditions. **A.** Histogram displaying low (0-6) moderate (6-16) and high (16+) number of division events for simulated cells under control conditions. Cells (4,000) were compiled across 4 drugs (0 dose), 100 cells per replicate, 10 replicates. **B.** Histogram displaying low (0-6) moderate (6-16) and high (16+) number of division events for simulated cells under trametinib treatment conditions (0.03 nM). Cells (1,000) were compiled across one dose, 100 cells per replicate, 10 replicates. **C.** Setup of the hypothesis, relating initial conditions of Generation 1 mother cells to the eventual number of divisions arising from them. **D.** Principal component plot of the initial states matrix under control conditions. Points are sized by number of division events, with colors equivalent to panel A. **E.** Top 20 loadings of the second principal component under control conditions, and a cartoon schematic of where they fall along the pathway driving the cell cycle. **F.** Projection of the trametinib data set onto the principal components

To answer these questions, we first focused on simulated control cells. The initial conditions of Gen 1 cells under control conditions were extracted into a cell-by-species matrix (4,000 simulated cells by 934 initial conditions, 4 control conditions, 100 cells for each, 10 replicates) (**Fig. 5C**). To identify species that may be associated with rapid division, we performed principal components analysis (PCA) of this matrix, and colored cells by their division phenotype (**Fig. 5D**). The second principal component (PC2) stratified Gen 1 mother cells based on the number of divisions. To identify the most important model species contributing to PC2, we analyzed the PCA loadings (**Fig. 5E**). Species with large loadings were associated with ERK signaling pathway components or downstream early cell cycle components. This suggests a simple hypothesis—that any fluctuation giving rise to higher ERK signaling capacity is associated with the rapid division phenotype under control conditions. Interestingly, PC1 did not correlate much with the rapidly dividing phenotype and was associated mainly with receptor-level species (**Fig. S2A**). A similar analysis approach, PLSR^86^, suggested the same as the PCA, although the first (not second) principal component was related to the rapidly dividing phenotype, and was also linked mainly to the ERK pathway signaling capacity (**Fig. S2B-C**).

Does this finding hold true under trametinib treatment conditions? That is, are cells with higher ERK signaling capacity more likely to retain rapid division phenotypes in the presence of sub-saturating doses of trametinib? Trametinib is a MEK inhibitor, and since MEK is a key component of the ERK signaling pathway, there is a clear connection. Trametinib treatment shifts the number of divisions distribution to the left, reducing the number of rapidly dividing cells (**Fig. 5B**). To answer the question, we generated a new initial Gen 1 mother cell state matrix from trametinib-treated simulated cells, and then applied the projection learned from PCA of the control data onto this matrix. The results of this projection indicate again that rapidly-dividing cells cluster towards higher values on PC2 (**Fig. 5F**). We conclude that in simulations, the rapidly dividing phenotype is driven by a multitude of factors impinging on higher ERK signaling pathway capacity, and these cells are likely to remain rapidly dividing in the presence of trametinib, contributing to acute resistance.

## Discussion

Data availability is a major bottleneck for systems biology model development. While there is a wide range of drug dose response viability assay data available, they are difficult to use for large-scale model development because simulation outputs often do not recapitulate the experiment outputs—cell number from cumulative division and death events in single cells. To address this gap, we developed a lineage-resolved simulation framework that tracks individual cell division and death events along with mechanistic detail that enables inference for why single cells have different outcomes. We demonstrate application of this framework using our previously developed model of proliferation and death signaling in single mammalian cells^53^, but in principle any model that simulates division and/or death events should be compatible, such as the one we present in addition^54^. We compare model simulations to experimental data for viability response to four different targeted anti-cancer drugs^55^. Discrepancies between model and data for palbociclib and neratinib, elaborated on further below, suggest where current understanding as captured by model assumptions is limited. Deeper analysis of trametinib cases suggest mechanisms of resistance and what drives rapidly cycling cells in general. Importantly, although we focus on four drugs here, previous work^61,62^ has reported similar data for 107 additional drugs studied in MCF10A, of which we believe approximately 65 that share targets with SPARCED model components are good candidates for similar modeling as here, which would be a logical next step.

For palbociclib, the simulations overpredicted its efficacy, showing very high growth inhibition at moderate doses to complete cytostasis at high doses. This reflects the indispensability of the drug target, CDK4/6, as per the model of the cell cycle pathway. However, in experiments, even the highest doses resulted in only partial growth inhibition. CDK4/6 is associated with traversing the cell cycle restriction point^87^. In pre-S-phase cells, one of the regulators of restriction point, Rb, is bound to a key transcription factor of the cell cycle process, E2F. In the presence of a growth stimulus, CDK4/6 is activated when bound to Cyclin D. This activated Cyclin D-CDK4/6 complex can phosphorylate and inactivate Rb, which then releases E2F. Subsequently, there is an upregulation of E2F which then mediates S-phase entry and progression by activating Cyclin E and Cyclin A. The mechanism of CDK4/6 inhibitors such as palbociclib attempt to induce cytostasis by preventing the inactivation of Rb by CDK4/6^88^. This canonical understanding places CDK4/6 as indispensable, similar to how it is modeled, but experiments did not agree with this assumption. One of the known resistance mechanisms of CDK4/6 inhibition is the loss of Rb function^89,90^. However, since MCF10A cells do not harbor such mutations, it is an unlikely explanation in this case. MCF10A are hormone receptor (HR) negative, whereas palbociclib is mainly understood to be more effective in HR positive contexts^91^, so perhaps differences caused by estrogen and progesterone receptor could be helpful in understanding the discrepancies. Another reported resistance mechanism in cancer cells is the overexpression of Cyclin E^92,93^, which is a regulator of the later stages of cell cycle, but is also not the case in MCF10A cells. In the results section, we investigated mismatch between model and experimental doubling time and restriction point behavior, finding neither likely to explain the discrepancies. Therefore, we think the most likely explanation is that CDK4/6 is simply not as indispensable for the cell cycle as contemporary views may portray. It has been increasingly reported that CDKs can compensate for one another^76^, so the activity of other CDKs could compensate for CDK4/6 activity in actively cycling cells. Such mechanisms were not included in the original cell cycle submodel^94^, so these additions are likely important for capturing effects of cell cycle-targeted therapies. Sensitivity analysis is a generally useful computational tool for understanding which mechanisms are related to particular data features, and may help such model refinement. However, currently, the proposed algorithm is computationally-intensive just for generating a single dose response data point (multiple replicates of 100s of initial stochastic cells). Increasing computational efficiency of the algorithm is an immediate next goal. Sensitivity analysis on stochastic models is notoriously difficult^95–97^ and an open area of research, but one that could synergize with approaches such as the one presented here.

For neratinib, the simulations underpredicted its efficacy, showing weak inhibition for moderate to high doses whereas the experiments showed significant growth inhibition to complete cytostasis within this range. To investigate this discrepancy, we considered the progression of ERK signaling within the single cell model and how the neratinib drug action might affect it. Neratinib is an irreversible inhibitor of EGFR, which attempts to block both ERK and Akt signaling by inhibiting ligand-receptor interactions. The model incorporates both ligand-induced and basal signaling along the ERK pathway. In simulations, if cells enter the cell cycle in the absence of ligand, it would result in proliferation that the drug action would be unable to inhibit. To test this, we performed subsequent simulations where EGF was absent from the growth media, but several simulated cells were still cycling. This is contrary to the experimental observations that MCF10A cells do not proliferate without EGF^82,83^, and explains the discrepancy observed between simulation and experimental results of neratinib dose response. Furthermore, we sought to account for this mismatch by altering a key model reaction modulating basal ERK activity, basal Ras-GDP to Ras-GTP exchange rate. We reduced this rate constant to minimize the probability of cell division in absence of EGF and ran all dose response simulations again. This time, the neratinib dose response showed closer alignment with the experimental result, but we observed overprediction of growth inhibition for all other drugs, presumably due to the altered balance between basal and ligand induced ERK signaling. Hence, ideally, the model should incorporate a more improved balance between basal and ligand induced signaling for describing cell proliferation events. In our previous work, stochastic single cell simulations initiated from a representation of a serum-starved MCF10A cell minimally entered S-phase without EGF^21^. However, for cell population simulations done here, single cells are subject to randomized sampling for induction of an asynchronously cycling population which more closely resembles the experimental conditions whereby drug treatment is applied after growth media is introduced to the cells. Also, cells were followed for much longer, which amplifies small percentages of cells still cycling with insulin treatment alone. Thus, the model’s limitations become more apparent here at the population level.

The neratinib case study highlighted an important future direction focused on parameter estimation for such models with stochastic components. This is a challenging area due to the computational cost of model evaluation, and the wide range of datasets that are needed to constrain large stochastic models. One part of our previous work was to do this for a subset of rate constants by “initialization”^21,98^. In initialization, certain model parameters and initial conditions are determined for a specific cell-line context using a set of focused parameter estimation operations which aim to tune parameters based on constraints placed on model observables. It is a computationally intensive process whereby each parameter estimation step performs iterative execution of deterministic model simulations. The SPARCED model is composed of a stochastic gene expression and a protein biochemistry module which are executed simultaneously. However, communication bottlenecks between the modules caused the computation time to be impractical for the purpose of initialization^98^. Recently, we solved the communication bottleneck problem which sped up the deterministic execution by over 200-fold^98^. Fast deterministic parameter estimation solvers have been reported for large-scale models as well^22^. This drastic increase in computation speed for deterministic simulations will allow a more exhaustive exploration of the model parameters essential for defining a more robust initialization protocol, but extending this to stochastic evaluation remains an important unsolved problem.

For trametinib, model predictions closely resemble experimental observations. Trametinib has high specificity for MEK1/2; once MEK1/2 is inhibited, it is no longer able to phosphorylate ERK1/2^99^. ERK signaling controls the G1/S-phase transition of the cell cycle^99^ through activation of RSK, which in turn upregulates the production of Cyclin D and CDK4/6. Cyclin D expression can drive the cell through the G1/S-phase checkpoint, as described above. When these events are inhibited by trametinib, the cell is unable to progress through the checkpoint. However, results indicate that low-to-medium doses of trametinib are unable to reduce cycling in every cell in the population. It was hypothesized that the signaling response patterns of a mother cell pass on to daughter cells, enabling an expression pattern to continue through multiple generations^100^. Therefore, we hypothesized that the initial values of the Generation 1 mother cells could predict the number of divisions that occur from said mother cell. The number of division events was explainable by principal components analysis, with higher ERK signaling capacity being associated with an increased number of division events. Thus, in this case, acute resistance to trametinib is simply related to a multitude of biochemical factors all impinging on increased activity of the target ERK pathway.

In conclusion, we have developed an algorithm that takes a mechanistically-detailed model of stochastic proliferation and death, and generates lineage-resolved simulations that can be used to interpret dose response viability data and better understand cellular response heterogeneity. Specific demonstrations suggested new insights into drug response, cell cycle biology, rapidly dividing phenotypes, and acute drug resistance. Given the extensive availability of drug dose response viability data, we anticipate that this work will help address the data availability bottleneck in modeling, facilitating the development of mechanistic models of single-cell behavior..

## Methods

### Code Availability

The final model scripts, files, and information are available on the SPARCED GitHub page at github.com/SPARCED/LinResSims. For detailed usage information, and for more details on how models and simulations are implemented, we recommend this page to the reader.

### Stochastic Model Components

The treatment of stochasticity was inherited from the SPARCED model and is described in detail there^21,53^. Briefly, stochasticity arises from gene expression, and it is described by what is sometimes called a telegraph model^101,102^. Genes in the model can be active or inactive, with first-order switching. Transcripts are produced from active genes and undergo degradation, also both first order. These reactions are simulated with a time step of 30 seconds, that was selected based on experimental data for gene switching rates in mammalian cells, to yield a low probability a gene becomes active and inactive in a single interval (in prior work faster time steps were confirmed not to impact simulation results). The reactions fire with a Gillespie/tau-leap-like mechanism. In a time step, random uniform numbers are compared to the gene activation and inactivation rate constants to determine gene switching events. The number of transcript births and deaths are determined by sampling from a Poisson distribution.

Heterogenization happens due to stochastic gene expression. At the time of cell division, the two daughter cells have identical molecule numbers and species concentrations. Thus, we default to a symmetric division. After cell division, stochastic gene expression happens in each cell independently, creating natural drift. It is in principle possible for a user to specify asymmetric division, which could be done by implementing a “divider” function^103^ which will be executed every time a division point is detected after any single cell simulation. Such a function may account for the individual protein molecule counts of the mother cell and determine their fate in the daughter cells as a result of an appropriate probabilistic operation.

In the example applied to the simple cell cycle model^54^ (see below), stochasticity was generated by creating an asynchronously cycling initial population as illustrated in Figure 1. That is, based on a time course simulation across a cell cycle, random time points were chosen, and the model state from these time points were used as initial conditions for different cells in the simulated population. The underlying model is deterministic and we found that its desired “cycling” behavior is very sensitive to parameter variation when we tried to generate stochasticity by adding random noise to the individual parameters and/or introducing Langevin equations to the model structure.

### SPARCED Pharmacodynamic Models

Reactions representing drugs binding to their reported targets with mass action rate laws were added to the SPARCED model (see model input text files). The assumptions and mechanism of action for each drug are described below. We tested each drug action model by observing simulated deterministic response of an average serum-starved cell to EGF and Insulin (growth media doses) with and without drug at high dose (10 μM). We required (i) intracellular and extracellular free drug concentration equilibrated rapidly (within a few minutes); (ii) drug-target engagement (complex formation) was observed similarly rapidly; (iii) that the drug had a substantial effect on a downstream biomarker (ppAKT-alpelisib, ppERK-trametinib, pEGFR-neratinib, or CDK4/6 activity-palbociclib).

*Alpelisib.* Alpelisib enters and leaves the cell with first-order kinetics and the same rate constant (0.01 s^-1^). Cytoplasmic alpelisib binds reversibly to its intracellular targets, p110 (representing all p110 isoforms) and free PI3K (p85/p110 heterodimers), with a dissociation constant (K_d_) of 2.4 nM^56^ and mass action kinetics (k_on_ = 0.001 nM^-1^s^-1^ ; k_off_ = 0.0024 s^-1^). Although alpelisib is a p110α isoform-specific inhibitor, the SPARCED model does not yet incorporate PI-3K isoform-specific biology, so this simplification is necessary at the current stage. The binding of alpelisib to p110 prevents its dimerization with the regulatory subunit (p85). Any drug-bound species loses its kinase activity. Any drug-bound species undergoes first-order degradation with a rate constant equal to that of the non-drug-bound species.

*Palbociclib.* Palbociclib enters and leaves the cell and the nucleus with first-order kinetics and the same rate constant (0.01 s^-1^). Nuclear palbociclib reversibly binds to its target, nuclear CDK4/6, with a dissociation constant (K_d_) of 1.9 nM and mass action kinetics (k_on_ = 0.001 nM^-1^s^-1^; k_off_ = 0.0019 s^-1^). Any drug-bound species loses its kinase activity. Any drug-bound species undergoes first-order degradation with a rate constant equal to that of the non-drug-bound species.

*Trametinib.* Trametinib enters and leaves the cell with first-order kinetics and the same rate constant (0.01 s^-1^). Cytoplasmic trametinib reversibly binds to its target, unphosphorylated free MEK, with a dissociation constant (K_d_) of 0.35 nM and mass action kinetics (k_on_ = 0.001 nM^-1^s^-1^ ; k_off_ = 0.00035 s^-1^). Any drug-bound species loses its kinase activity and ability to bind substrates. This is a simplification of trametinib action but is effective for capturing the broad effects of downregulating the ERK pathway. Any drug-bound species undergoes first-order degradation with a rate constant equal to that of the non-drug-bound species.

*Neratinib.* Neratinib enters and leaves the cell with first-order kinetics and the same rate constant (0.01 s^-1^). Cytoplasmic neratinib binds irreversibly to free EGFR, ErbB2, and ErbB4 with first-order kinetics (k_on_ = 10^-4^ nM^-1^s^-1^). While this is a kinase inhibitor, and drug bound complex loses kinase activity, for simplicity we disallow subsequent interaction with other receptors and ligands. Any drug-bound species undergoes first-order degradation with a rate constant equal to that of the non-drug-bound species.

### Lineage-Resolved Simulations

#### Asynchronous population

Cell population simulations are initiated by creating a representation of an asynchronously cycling cell population. The starting size of the cell population is specified by the user. For each starting cell, initial conditions representing an average serum-starved MCF10A cell are used to create a heterogenized cell population^21^. For heterogenization, we run stochastic single cell simulations for 48 simulated hours under serum-starved conditions, using the initial conditions of the average serum-starved MCF10A cell. Thus, the intrinsic gene expression noise incorporated within the single cell model leads to heterogeneity in protein levels across the starting cell population over the duration of simulation time. Then, simulated growth media with EGF (3.3 nM) and insulin (1721 nM) is introduced and another series of stochastic simulations are run for each individual cell for 48 hours. From the generated trajectories, for each cell a timepoint is randomly selected from a uniform distribution using the NumPy `randint` function. The conditions at this time point for each cell are used as the initial conditions for the first generation. Single-cell simulations are executed for all first generation cells for the user-specified duration (typically 72 hours).

#### Identifying cell division events

Once the single-cell simulations are completed, the generated outputs are analyzed to determine cell division events. The cell division events are detected by analyzing Cyclin B-CDK1 trajectories. For this, we defined a python function combining the `find_peaks` methods in the SciPy signal processing library and the n-th discrete difference calculation method (along any given axis) in the NumPy library. For any individual cell, if a division event is detected, timepoints after the occurrence of cell division events are discarded and the state vector at the time of cell division is selected as the initial condition for two new second generation cells. Thus, we assume symmetric division, where the daughter cells have identical initial conditions. Importantly, daughter cells immediately begin experiencing drift from one another due to stochastic gene expression, which is constantly occurring in every simulated cell differently.

#### Identifying cell death events

Cleaved PARP is the readout for cell death^21^. For any single cell, if more than half of PARP has been cleaved at any time point, the cell is labeled dead at that time point. To compare simulated cell death events to experimental data, we assumed that any death event within the last 1 hour of the simulation would be observable by the viability staining used.

#### Subsequent generations

For each generation, state matrices for individual cells are obtained and saved as part of output dataset. In the event of a cell division, we retain the state matrix of the mother cell until the time point of division and the remaining portion is truncated and discarded. For every cell, we scan the output for the duration of its lifetime to find division events. To determine the required simulation time for next generation of daughter cells, the division time point is subtracted from the total simulation time. The single cell outputs at the time point of division of each mother cell is recorded as initial conditions for the next generation of daughter cells. Thus, we define the required simulation time, population, and initial conditions for the next generation of cells. This process is repeated for the subsequent generations of cell populations. In a given generation, if there is no cell division event observed within the simulation time, the population simulation is terminated. We assume symmetric division, where the daughter cells have identical initial conditions.

#### Implementation

Computation is performed using HPC-compatible parallel processing in Python whereby single cell simulations are run in individual CPU threads. To run the cell population simulation, a computational environment with an implementation of MPI (Message Passing Interface), such as OpenMPI^104^ on Linux and MSMPI on Windows systems needs to be set up in addition to the dependencies of the SPARCED model pipeline. Before the simulation can be performed, the SPARCED model is built using the python script under scripts/createModel.py, which creates an executable single cell model based on the specifications in the input files. Once the model build process is complete, MPI is be used to run cell population simulations (see git repository). Upon completion of simulations, the results are saved to disk as Python pickle objects for analysis and visualization. For detailed reproduction of results in the paper and for specific use of the codebase, we would refer the reader to the GitHub documentation.

A main function is the **`cellpop.py`** script. The command line arguments passed to the this script are used to specify inputs to the simulation representing the experimental conditions as well as several workflow parameters that dictate the computation. A more detailed specification of these variables can be made by using a json config file, which the user may define for each simulation run. This allows the alteration of several key workflow parameters without modification of the simulation script itself. By default, simulation config files are located in the folder sim_configs and each file is passed to the cellpop python script using the argument –sim_config. The contents of the sim_config file are read as a python dict object by the **`cellpop.py`** script. We refer the reader to the GitHub repository for detailed usage of the config file.

### Running Cell Population Simulations with a New Single Cell Model

By default, the cell population simulation workflow uses the SPARCED single cell model. It is capable of running simulations with a different single cell model given that the model has a compatible structure. We have provided an example applied to a simple, classical cell cycle model^54^ (see the GitHub repository), but others could be compatible. A compatible model must (i) have a state matrix representing a single cell, (ii) have a variable representing the dynamic molecular signature of cell cycle markers, i.e., periodic activation and inactivation of cyclins, and (iii) be executable within a python module.

To replace the SPARCED model in cell population simulations with another single cell model:

1. Place all single cell simulation operations within a python function. This function must be given a unique name and saved in a module of the same name under bin/modules. The name of the module must be specified under “model_module/run_model” option in the config file. This function may accept any number of arguments required to execute the single cell model (e.g. initial conditions, parameters, duration etc.) and the arguments must be correctly mapped within the “kwargs_default” dictionary in the next step.
2. Write another python function to generate an input dict for the single cell model function, mirroring the input/output structure of the LoadSPARCED function. This function should be given a unique name and saved in a module of the same name as the function under bin/modules. The name of the module must be specified under “model_module/load_model” option in the config file. The function must return two dictionaries, namely:

- model_specs: dictionary containing “species_all” (list of model species names according to their order in the state matrix) and “cc_marker” (name of the cell cycle marker species)
- kwargs_default: dictionary containing keyword arguments for the “run_model” function.
3. Save both python functions as modules with the same name as the functions under bin/modules.
4. Write a json config file with key-specific values appropriate for the new model structure. Be sure to make “load_model” and “run_model” options consistent with the new module names. For more details on the structure of the sim config, see sim_configs/README.md

As an example for this procedure, we used a classic simple ODE model of the cell cycle^54^. The model represents the interactions of cdc2 and cyclin during major events of the cell cycle in Xenopus oocytes. This particular implementation of the model has been defined entirely within a python module (bin/modules/TysonModule.py) and simulated using the LSODA solver in the `scipy` library. The periodic oscillation of cyclin-P/Cdc2 complex has been selected as the cell cycle marker in this implementation. The “load_model” and “run_model” modules have been provided as bin/modules/LoadTyson.py and bin/modules/RunTyson.py. The sim_config json file corresponding to this workflow is sim_config/default.json.

### GR Score Calculation and Experimental Data Source

Experimental data^55^ were obtained from synapse (www.synapse.org/Synapse:syn18456348/wiki/590585), and the data pull script is provided in the GitHub. Dose responses were calculated using the growth rate inhibition metric (GR)^60^. Dose response simulations were run for 10 dose-levels matching experimental data for each drug and 10 replicates of each dose. Outputs from the cell population simulations were read and analyzed to determine the total number of living cells over time for the duration of the experiment time. The GR scores were computed for each replicate from the number of living cells at 72 hours using the Python script provided as part of GR-metrics git repository.

### Calculation of Fractional Cell Cycle Progression

#### Cell cycle progress estimation

For the palbociclib dose response, the extent of cell cycle progress at the time of drug addition was estimated using a function of average cyclin concentration levels. CyclinE-CDK2, CyclinA-CDK2 and CyclinB-CDK1 species concentrations were converted to a relative measure based on their observed peaks. An average of these three variables over time generated an oscillating function, with trough-to-trough distance representing total cell cycle time. We determined an average trajectory of this function using a deterministic simulation. Then, we calculated the function trajectory for individual Gen 1 cells. We calculated the relative cell cycle progression as cell cycle progression time as aligned to the average, divided by the time between two neighboring troughs in that cell’s trajectory.

#### Estimation of cell division probability given cell cycle progression

For any given drug dose, the cell cycle progression of all cells at the time of dose administration was calculated. Then all living cells were grouped into those that divided and did not divide in both Gen 1 and Gen 2. Gaussian kernel density estimation was used to estimate the probability density function for each group. Using the probability density function, the number of cells for dividing and non-dividing groups within cell cycle progress time intervals with increments of 0.01 were estimated. For each interval, the probability of division at Gen 1 and 2 were calculated using the ratio of number of dividing cells and total number of cells.

### Principal Components and Partial Least Squares Analysis

The number of progeny arising from each Gen 1 cell (‘mother cell’) was determined from control condition simulations as above. Z-score normalization was applied to the initial condition matrix (cells-by-species) using `standardScaler.transform` in scikit-learn. Principal component analysis was completed using `decompostion.PCA` in scikit-learn. To apply this projection to the drug-treated simulated cells, we generated a new initial state matrix from simulated mother cells treated with 0.032 nM trametinib and normalized as above. Partial Least Squares Regression was completed using the cross_`decomposition.PLSRegression` package in scikit-learn. This was applied to the initial values matrix of control condition simulations and weights extracted from the model object using the .x_loadings attribute. The code underlying this analysis is in the above-mentioned GitHub repository as well.

## Supporting Information Legends

**Figure S1.**
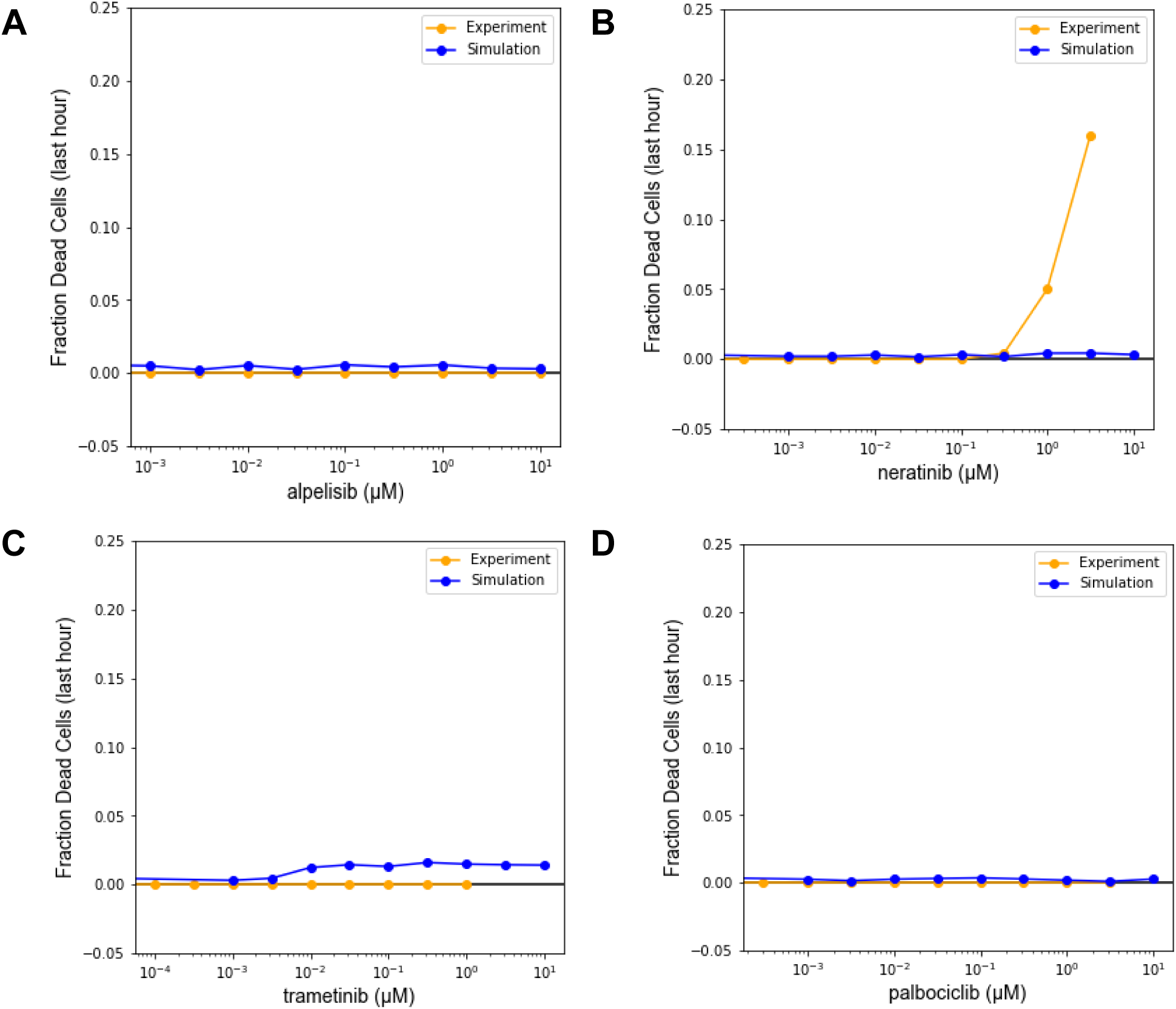
Cell Death Analysis. **A-D.** Fraction of dead cells among cells simulated between the timepoints 71h and 72h (the last hour) for the four drugs as indicated. Experimental data were obtained as indicated in the main text references.

**Figure S2.**
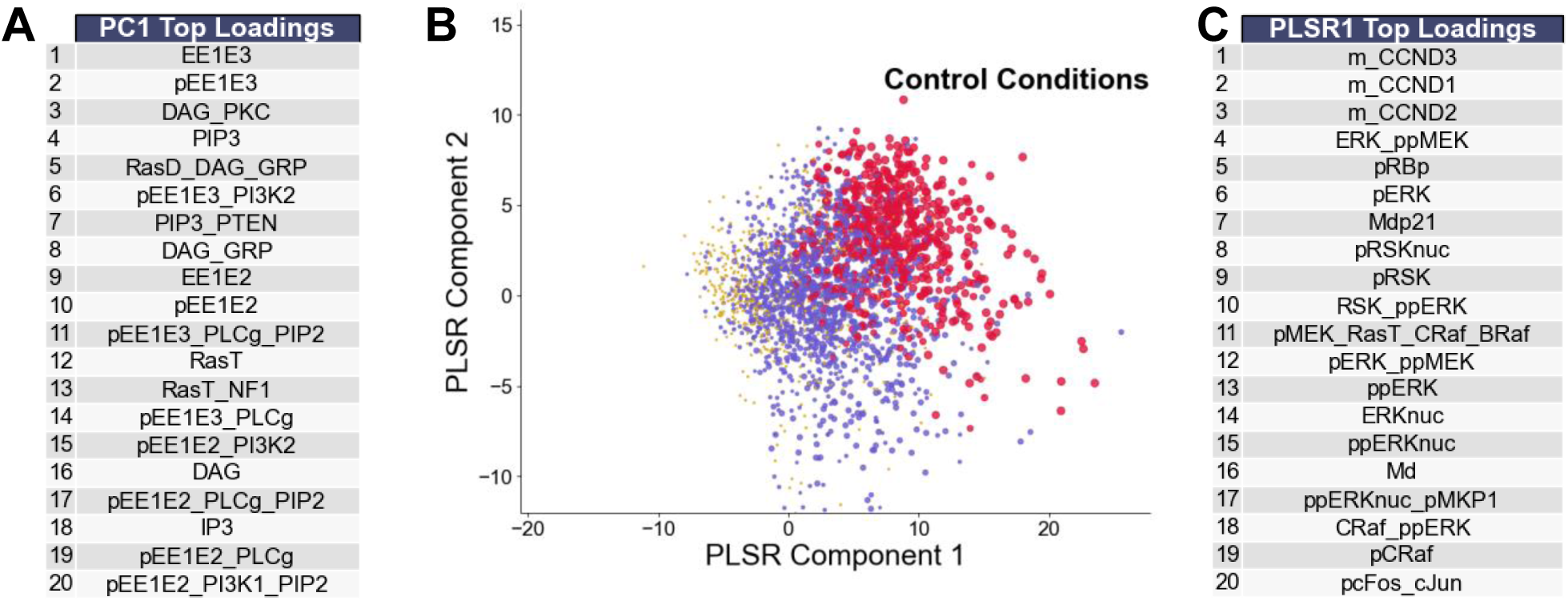
Rapidly Dividing Phenotype Analysis. Initial conditions from 4,000 simulated cells under control conditions were arranged into a matrix as in Figure 5 and analyzed by principal components analysis (PCA) or partial least squares regression (PLSR). **A.** Top 20 loadings of the first principal component under control conditions. **B.** PLSR analysis. Points are sized by number of division events, with colors equivalent to Figure 5A. **C.** Top 20 loadings of the first PLSR component under control conditions.

## Acknowledgements

We thank Michael Salim, Nicole Hobbs, and Daniel Cook for helpful discussions. This work was supported by NIH grant R35GM141891 to MRB. The funders had no role in study design, data collection and analysis, decision to publish, or preparation of the manuscript. We thank the Clemson University Palmetto team for use of high-performance computing resources, which is supported by the National Science Foundation under Grant Nos. MRI# 2024205, MRI# 1725573, and CRI# 2010270. Any opinions, findings, and conclusions or recommendations expressed in this material are those of the author(s) and do not necessarily reflect the views of the National Science Foundation. We also acknowledge the Umea University high-performance computing cluster.

## CRediT Author Contributions

Arnab Mutsuddy: Data Curation, Formal Analysis, Investigation, Methodology, Software, Validation, Visualization, Writing – Original Draft Preparation, Writing – Review & Editing Jonah Huggins: Data Curation, Formal Analysis, Investigation, Methodology, Software, Validation, Visualization, Writing – Original Draft Preparation, Writing – Review & Editing Aurore Amrit: Data Curation, Formal Analysis, Investigation, Methodology, Software, Validation, Visualization, Writing – Original Draft Preparation, Writing – Review & Editing Atalanta Gasaway: Software, Validation Cemal Erdem: Supervision, Writing – Review & Editing Jon Calhoun: Supervision Marc Birtwistle: Conceptualization, Funding Acquisition, Project Administration, Supervision, Writing – Original Draft Preparation, Writing – Review & Editing

